# Microglia regulate early postnatal development of corticolimbic GABAergic synapses and anxiety-related behavior

**DOI:** 10.64898/2026.11.23.768483

**Authors:** Federico Rotondo, Kyra Sohns, Tamara C. Ritter, Juliette Soubra, Michael J. Schmeisser, Ari Waisman, Dilja Krueger-Burg

## Abstract

Microglia have emerged as key regulators of synaptic remodeling, including GABAergic synapses, but their contribution to the development of anxiety-related circuits in the corticolimbic system remains poorly understood. Here, we investigated the consequences of postnatal microglial depletion on GABAergic synapses, inhibitory synaptic transmission, and anxiety-related behaviors in mice. Microglia were pharmacologically depleted during either the first (P1–P14) or second (P15–P28) two postnatal weeks. Early, but not late, postnatal microglial depletion reduced the number of perisomatic GABAergic synapses in the amygdala and increased the number of gephyrin puncta across several corticolimbic regions. Furthermore, early microglial depletion disrupted the development of normal anxiety-related behaviors. These effects were independent of microglial GABA_B1_ receptor expression, suggesting that GABA-receptive microglia do not play a significant role in these processes. Together, our findings support the notion that microglial remodeling of corticolimbic GABAergic synapses is required for the physiological development of anxiety-related circuits, providing a basis for future studies on the implications for anxiety disorders.

## Introduction

Information flow throughout the forebrain is critically shaped by GABAergic inhibitory circuits and synapses, which play a fundamental role in regulating neuronal excitability, governing the temporal and spatial dynamics of neuronal activity, and orchestrating neuronal ensembles to coordinate network integration (*1, 2*). Accordingly, GABAergic dysfunction and an imbalance in the excitation/inhibition ratio have been implicated in several psychiatric and neurodevelopmental disorders (*3–9*), and GABAergic synaptic transmission has emerged as an important target for psychiatric pharmacotherapies (*10–12*). A particular vulnerability of the GABAergic system is observed in anxiety disorders (*7, 8, 13*), which are among the most prevalent mental health conditions worldwide and which place a substantial burden on affected individuals, their families, and society (*14*). Defining the mechanisms that contribute to the development and function of GABAergic circuits and synapses that encode anxiety-related behaviors is therefore critical for understanding the emergence of these disorders.

In recent years, increasing evidence has highlighted microglia as key players in shaping the development and maintenance of neuronal circuits and synapses. Microglia are the resident macrophages of the central nervous system, and their role was originally assumed to consist primarily in protecting the brain against the consequences of infection and tissue damage (*15, 16*). It has long since been recognized, however, that microglia also contribute to a number of essential brain functions under physiological, homeostatic conditions, including the regulation of synaptic structure and function. Initially, it was demonstrated that microglia are instrumental in the pruning of synapses via the phagocytosis of synaptic material (*17*), and that this process occurs under a wide range of physiological conditions, both during early postnatal development and in adulthood during learning and memory formation (*18, 19*). More recently, it was found that they also contribute to initial synaptogenesis and maintenance of synapses, leading to the general concept of synaptic remodeling as a key function of microglia (*20–22*). While the vast majority of research has focused on their interaction with glutamatergic excitatory synapses, a smaller number of recent studies have demonstrated that microglial remodeling of GABAergic synapses is likely to be equally important (*23, 24*) and may be mediated by a subset of GABA-receptive microglia (*25, 26*). Given that microglia are known to contribute to the pathogenesis of neuropsychiatric disorders, including anxiety disorders (*27–29*), it is plausible that this link may involve their effects on GABAergic circuits. To date, however, virtually no information is available on the role of microglia in shaping GABAergic synapse development in the brain regions and circuits that are known to encode anxiety behaviors and related pathologies, most notably including the corticolimbic system (*30–32*).

In the present study, we sought to address this issue by investigating the consequences of manipulating microglial function in mice on postnatal development of GABAergic synapses in three interconnected regions of the corticolimbic system, the medial prefrontal cortex, the amygdala, and the ventral hippocampus, all of which are closely linked to anxiety disorders and are prominently regulated by GABAergic inhibition (*8, 33–35*). To this end, microglia were pharmacologically depleted via administration of the CSF1R inhibitor PLX5622 either in the first two postnatal weeks (P1-14), which mark the peak period of synaptogenesis, or in the second two postnatal weeks (P15-P28), which represent a period of activity-dependent synaptic refinement and synapse elimination (*36–38*). Using a combination of immunohistochemical, electrophysiological and mouse behavioral approaches, we assessed the resulting effects on GABAergic synaptic markers, including the presynaptic vesicular inhibitory amino acid transporter (VIAAT) and the postsynaptic scaffolding protein gephyrin, as well as on inhibitory synaptic transmission and on behavioral measures of anxiety and fear-related learning. In addition, we investigated the role of GABA-receptive microglia in these processes using microglia-specific deletion of the GABA_B1_ receptor. Our findings support the notion that microglial remodeling of corticolimbic GABAergic synapses is indeed required for the physiological development of anxiety-related circuits, providing a basis for future studies on the implications for anxiety disorders.

## Results

### Microglia interact differently with inhibitory synapses across critical developmental periods

In order to investigate whether microglia may play a role in the developmental remodeling of GABAergic inhibitory synapses in the corticolimbic system, we first sought to define the interaction between them across different developmental timepoints and corticolimbic regions (Figure 1A-I). It has previously been shown that the pruning of GABAergic synapses by microglia can be detected by assessing the quantity of GABAergic synaptic markers within microglial lysosomes (*25*). Using an immunohistochemical analysis (Figure 1A-C) for the microglial marker IBA1 (ionized calcium-binding adaptor molecule 1), the microglial lysosomal marker CD68, and the GABAergic presynaptic protein VIAAT (vesicular inhibitory amino acid transporter), we quantified the colocalization between CD68 and VIAAT across three brain regions (medial prefrontal cortex, mPFC; basolateral amygdala, BLA; and ventral hippocampus, vHPC; Figure 1D-F). We focused on four developmental timepoints (postnatal days P7, P14, P21, and P28), which span the peak periods of initial synaptogenesis and subsequent activity-dependent synapse refinement in the mouse forebrain (Figure 1G). Analysis of the percentage of CD68^+^ lysosomes colocalizing with VIAAT (Figure 1H) indicated that between 10-40% of microglial lysosomes contained presynaptic material from GABAergic synapses in all three brain regions, with a trend toward a reduction across developmental timepoints (Figure 1I-K). Conversely, analysis of the percentage of VIAAT puncta colocalizing with CD68 (Figure 1L) revealed that only a small fraction of VIAAT-containing synaptic structures (< 5%) is actively being pruned by microglia across all brain regions and timepoints (Figure 1M-O). These findings indicate that a substantial proportion of microglia actively interact with GABAergic synapses across developmental timepoints in the corticolimbic system, but that the specific contribution of microglial pruning to the overall GABAergic synaptic landscape may be relatively minor.

**Figure 1.**
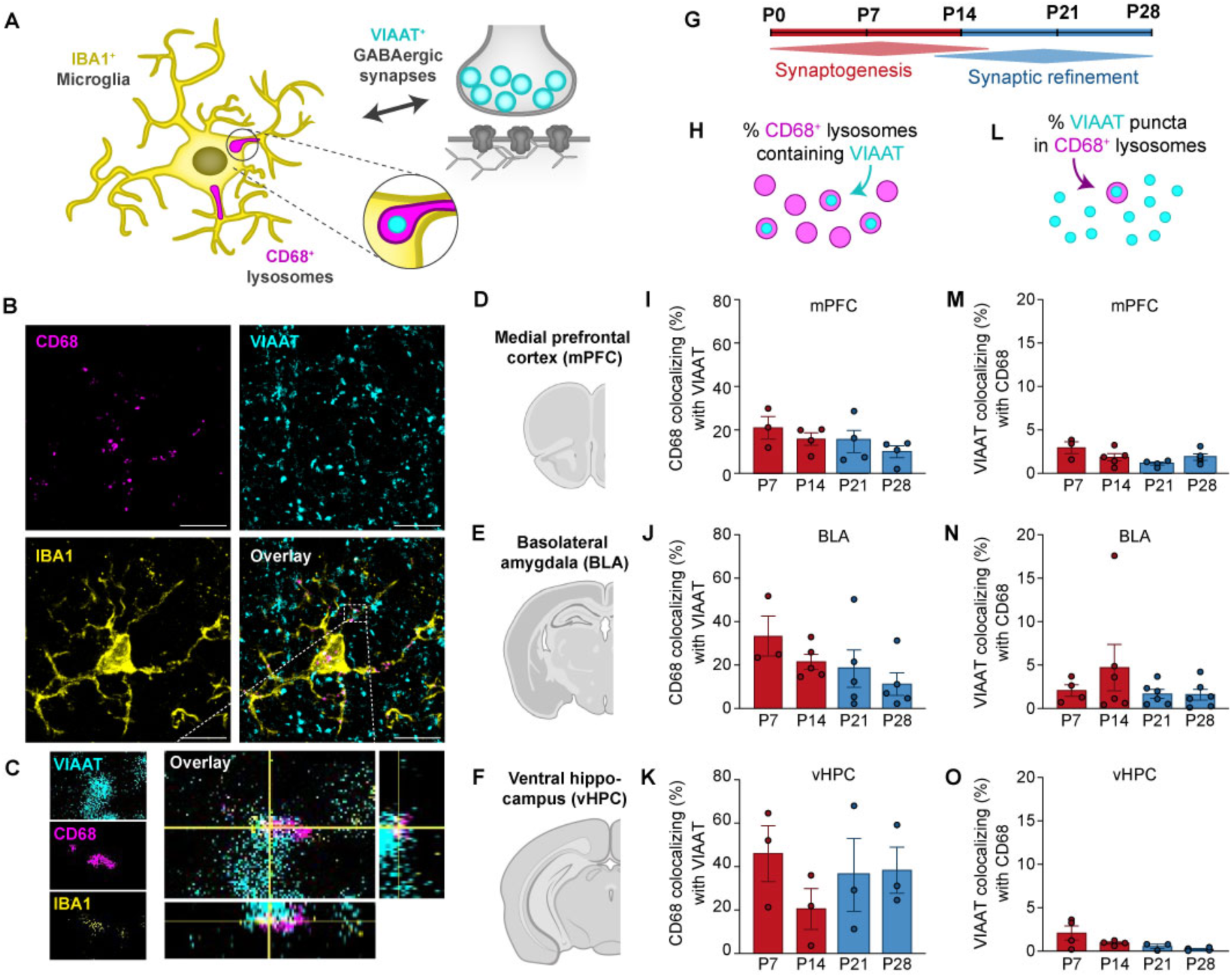
Microglia interact with GABAergic synapses across critical developmental periods in the corticolimbic system. (**A**) Schematic representation of the interaction between microglia and GABAergic synapses (labeled with IBA1 and VIAAT immunohistochemistry, respectively), including microglial lysosomes (labeled with CD68 immunohistochemistry). (**B**) Representative photomicrograph of immunostaining for IBA1 (yellow), CD68 (magenta) and VIAAT (cyan). (**C**) Magnified image of an IBA1- and CD68-positive microglial lysosome containing VIAAT. Right: YZ orthogonal view of CD68 punctum colocalizing with VIAAT; bottom: XZ view. (**D-F**) Anatomical localization of brain regions analysed, including mPFC (D), BLA (E), and vHPC (F). (**G**) Developmental timeline of synapse formation and refinement in the mouse corticolimbic system. (**H**) Schematic representation of the quantification of the percentage of CD68-positive lysosomes containing VIAAT. (**I-K**) Quantification of the percentage of CD68-positive lysosomes containing VIAAT puncta at P7, P14, P21 and P28 in mPFC (I), BLA (J), and vHPC (K). (**L**) Schematic representation of the quantification of the percentage of VIAAT puncta found in CD68-positive lysosomes. (**M-O**) Quantification of the percentage of CD68-positive lysosomes containing gephyrin puncta at P7, P14, P21 and P28 in mPFC (M), BLA (N), and vHPC (O). mPFC P7, n = 3; mPFC P14-28, n = 5; BLA P7, n = 4; BLA P14-28, n = 5-6; vHPC P7-14, n = 4; vHPC P21-28, n = 5. Error bars represent SEM.

### Absence of microglia during early postnatal development impacts GABAergic synapses differently in the BLA and IL

To further define the function of microglia in shaping corticolimbic GABAergic synapses, we next asked whether the absence of microglia during the first two weeks of life, a period of intense synaptogenic activity (*17, 39, 40*), might alter the number of GABAergic synapses in these regions. To this end, we administered PLX5622, a colony stimulating factor 1 receptor inhibitory which selectively deplete microglia from the brain (*41*), daily to newborn pups from P1-P14 (Fig. 2A, Fig. S1A), resulting in a near-complete depletion of microglia from the entire brain (Fig. 2B-C, Fig. S1B). Using an immunohistochemical approach, we subsequently determined whether GABAergic synapses were altered in the ventral hippocampus (area CA1, layers stratum pyramidale, SP, and stratum radiatum, SR), amygdala (basolateral amygdala, BLA, and central amygdala, CeA) and medial prefrontal cortex (prelimibic cortex layer II/III, PL, and infralimbic cortex layer II/III, IL) (Fig. 2D). Interestingly, daily application of PLX5622 slightly reduced the growth rate of pups during the first two weeks (Fig. S1C), suggesting side effects of CSF1R inhibition of myeloid cells on physical development (*42*).

**Figure 2.**
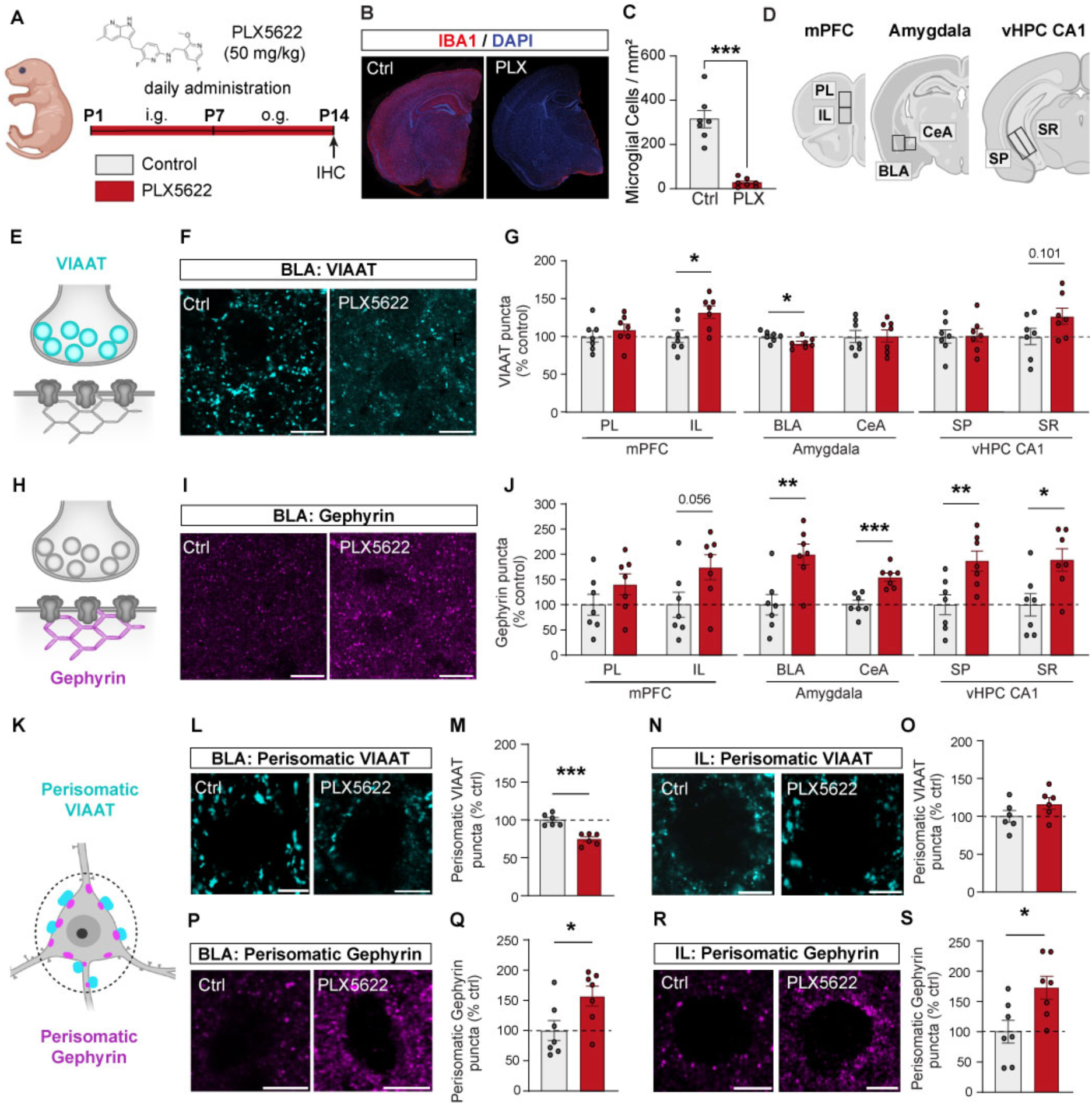
Absence of microglia from P1-P15 impacts GABAergic synapses differently in the BLA and IL. (**A**) Experimental design for PLX5622-mediated depletion of microglia from P1-P14 and immunohistochemical analysis at P14. Image of mouse generated using Biorender. (**B**) Representative photomicrographs of IBA1 immunostaining in control and PLX5622-treated mice. (**C**) Quantification of microglial cell density in control and PLX5622-treated mice (n = 7). (**D**) Anatomical localization of brain regions analysed. **(E)** Synaptic localization of pre-synaptic marker VIAAT. (**F**) Representative photomicrograph of VIAAT immunostaining in control and PLX-treated conditions in the BLA. Scale bar 10 μm. (**G**) Quantification of the number of VIAAT puncta in control (light grey) and PLX5622-treated (red) mice in each brain region investigated, expressed as percentage of control. (**H**) Synaptic localization of post-synaptic marker gephyrin. (**I**) Representative photomicrograph of gephyrin immunostaining in control and PLX-treated conditions in the BLA. Scale bar 10 μm. (**J**) Quantification of the number of gephyrin puncta in control (light grey) and PLX5622-treated (red) mice in each brain region investigated, expressed as percentage of control. (**K**) Schematic representation of the immunohistochemical analysis of perisomatic puncta. Presynaptic VIAAT puncta in the perisomatic area and axon initial segment are shown in cyan, while postsynaptic gephyrin puncta are shown in magenta. (**L**) Representative photomicrograph of perisomatic VIAAT immunostaining in control and PLX5622-treated conditions in the BLA. Scale bar 2 μm. (**M**) Quantification of the number of perisomatic VIAAT puncta in the BLA, expressed as percentage of control (n = 6). (**N**) Representative photomicrograph of perisomatic VIAAT immunostaining in control and PLX5622-treated conditions in the IL. Scale bar 2 μm. (**O**) Quantification of the number of perisomatic VIAAT puncta in the IL, expressed as percentage of control (n = 6). (**P**) Representative photomicrograph of perisomatic gephyrin immunostaining in control and PLX5622-treated conditions in the BLA. Scale bar 2 μm. (**Q**) Quantification of the number of perisomatic gephyrin puncta in the BLA, expressed as percentage of control (n = 7). (**R**) Representative photomicrograph of perisomatic gephyrin immunostaining in control and PLX5622-treated conditions in the IL. Scale bar 2 μm. (**S**) Quantification of the number of perisomatic gephyrin puncta in the IL, expressed as percentage of control (n = 7). Error bars represent SEM. Statistically significant comparisons are marked with * p < 0.05, ** p < 0.01, *** p < 0.001.

However, this weight reduction was reversed once PLX5622 administration was discontinued, with normal body weights observed in adulthood (Fig. S1D). Analysis of the presynaptic vesicle protein VIAAT (Fig. 2E-G) revealed a small but significant decrease in the number of VIAAT puncta in the BLA (9% compared to control), a significant increase in the IL (32% compared to control), and no significant alterations in any other brain region analysed (Fig. 2G). These findings indicate that microglia differentially affect GABAergic presynapses during the period of synaptogenesis in a brain region-dependent manner. Surprisingly, investigation of the postsynaptic scaffolding protein gephyrin (Fig. 2H-J) revealed a substantial and significant increase in the number of gephyrin puncta in almost all the regions analysed (Fig. 2J). Similar increases, albeit smaller in magnitude, were found in a subset of regions for the postsynaptic adhesion protein Neuroligin-2 (Nlgn2), but not for the GABA_A_Rγ2 subunit, a core subunit of the ionotropic GABA_A_ receptor (Fig. S2A-F). These data indicate that GABAergic pre- and postsynapses are differentially regulated by microglia, and that the absence of microglia induces a pronounced upregulation in GABAergic postsynaptic scaffolding components independently of the presence of corresponding presynaptic terminals.

### Absence of microglia during early postnatal development differently impacts perisomatic GABAergic synapses

The GABAergic system is highly diverse, with different GABAergic neuron subtypes forming synapses in different subcellular locations of their target neurons. Among others, parvalbumin-positive (PV^+^) neurons specifically form synapses onto the perisomatic region and axon initial segment of target neurons, and PV^+^ synapses were previously shown to be modulated by microglial function in the somatosensory cortex (*25, 26*). To determine whether the changes observed previously (Fig. 2I, M) may result from effects on PV^+^ perisomatic synapses, we analysed synaptic puncta selectively in the perisomatic area of cells in the BLA and IL (Fig. 2K). Interestingly, in the BLA, the analysis of perisomatic VIAAT puncta (Fig. 2L-M) revealed a more prominent decrease (25.4%) compared to the one observed in the general analysis (Fig. 2G, BLA), suggesting that the absence of microglia induces a stronger effect on GABAergic synapses in the perisomatic area compared to the other subcellular locations of the target neurons. In contrast, analysis of VIAAT in the perisomatic area of the IL (Fig. 2 N-O) revealed a trend but no significant difference between PLX5622 and the control group, indicating that in IL, perisomatic presynapses may be less affected by microglial depletion than those in other subcellular locations. Analysis of the perisomatic gephyrin puncta in the BLA (Fig. 2P-Q) and IL (Fig. 2R-S) revealed an analogous increase compared to the one observed in the general analysis (Fig. 2I, BLA and IL), indicating that these changes in gephyrin puncta do not differentially affect synapses in different subcellular locations. Together, these findings indicate that depletion of microglia during synaptogenesis differentially affects presynaptic terminals but not postsynaptic scaffolding structures at perisomatic vs. other subcellular locations. Whether these changes are specific to PV^+^ synapses or may also affect other GABAergic perisomatic synapses remains to be determined.

### Absence of microglia during early postnatal development impacts synaptic transmission at GABAergic synapses in the BLA but not in the IL

To further investigate whether the changes in GABAergic pre- and postsynaptic marker proteins observed in the BLA and IL are mirrored by corresponding alterations in inhibitory synaptic transmission, we performed patch-clamp recordings from acute slices prepared from P14-P16 pups following PLX5622 administration (Fig. 3A-K). In accordance with the previously observed immunohistochemical alterations, we found a significant reduction of the frequency of miniature inhibitory postsynaptic currents (mIPSCs) in the BLA (Fig. 3F), as well as an increase in decay time (Fig. 3I). No alterations were recorded in the amplitude (Fig. 3G) and rise time of mIPSCs (Fig. 3H). These alterations, together with the reduction in VIAAT puncta (Fig 2G, M) are consistent with the notion that GABAergic presynaptic terminals are reduced in the BLA in response to microglia depletion during synaptogenesis. In contrast, analysis of mIPSCs in the IL revealed no difference between PLX5622-treated animals and the control group (Fig. 3J-M), matching the data on VIAAT puncta in the perisomatic region (Fig. 2O) but not on general VIAAT puncta (Fig. 2G). Given that mIPSC recordings are known to more prominently reflect synaptic transmission at perisomatic compared to dendritic synapses (*43, 44*), these findings are consistent with the notion that microglial depletion may result in a specific increase in GABAergic presynaptic terminals onto dendritic but not perisomatic subcellular locations in the IL.

**Figure 3.**
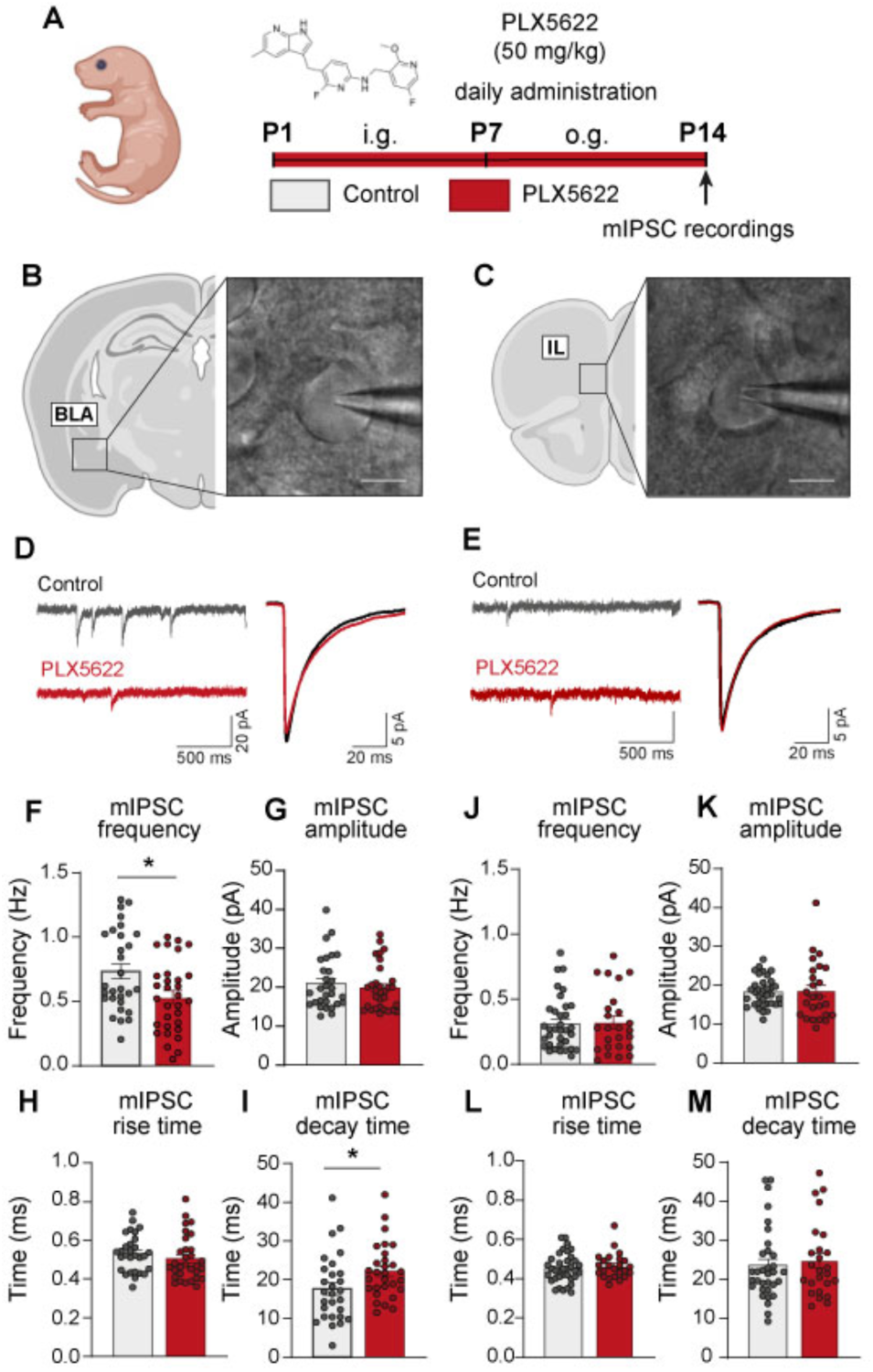
Absence of microglia from P1-P14 impairs synaptic transmission at GABAergic synapses in the BLA but not the IL. **(A)** Experimental design for PLX5622-mediated depletion of microglia from P1-P14 and electrophysiological analysis at P14-P16. Image of mouse generated using Biorender. **(B-C)** Representative image of whole-cell patch configuration in the BLA (B) and IL (C). **(D-E)** Representative recording traces from control (light grey) and PLX5622-treated (red) mice in the BLA (D) and IL (E). **(F-I)** Quantification of mIPSC frequency (F), amplitude (G), rise time (H), and decay time (I) in control (light grey) and PLX5622-treated (red) mice in the BLA (5 mice per condition; frequency, n = 29, 31 cells; amplitude, n = 29, 31 cells; rise time, n = 23, 25 cells; decay time, n = 28, 29 cells). **(J-M)** Quantification of mIPSC frequency (J), amplitude (K), rise time (L), and decay time (M) in control (light grey) and PLX5622-treated (red) mice in the IL (5 mice per condition; frequency, n = 34, 25 cells; amplitude, n = 35, 25 cells; rise time, n = 36, 24 cells; decay time, n = 33, 25 cells). Error bars represent SEM. Statistically significant comparisons are marked with * p < 0.05.

### Absence of microglia during early postnatal development results in altered anxiety-related behaviors in adulthood

Given that GABAergic synapses and circuits in both the BLA and the IL are known to play key roles in anxiety- and fear-related neuronal circuits, we next investigated whether microglia depletion during synaptogenesis results in long-term alterations in defensive behaviours in adulthood. To this end, mice received administration of PLX5622 from P1-P14 as previously described, and a panel of behavioral tests was conducted at 8-10 weeks of age (Fig 4A), including the elevated plus maze (EPM), open field test (OFT) and associative fear conditioning test (AFC) (Fig. 4B). In the EPM paradigm (Fig. 4C), in which mice explore an apparatus consisting of two open arms, deemed anxiogenic, and two closed arms, considered to be safe areas, control mice spent significantly less time in the open arms than the closed arms. This avoidance of the open arms indicates that control mice correctly identified open spaces as a more threatening, anxiety-inducing environment. In contrast, PLX5622-treated mice interestingly made no distinction between open and closed arms, spending equal time in both arms, which indicates that normal threat perception and anxiety-related risk assessment were disrupted following microglial depletion (Fig. 4D-E). Importantly, no PLX5622-related alterations in total locomotor activity were recorded on the EPM, confirming that microglial depletion did not cause non-specific effects on locomotion (Fig. 4F). Following the EPM, mice were tested in an OFT paradigm, in which mice are permitted to explore a novel arena which includes an open center, considered to be anxiogenic, and a periphery that is considered to be a safe area (Fig. 4G). Unlike in the EPM, no differences were observed between control and PLX5622-treated mice in the time spent in the center of the OFT setup or the periphery (Fig. 4H-I). Moreover, no alterations in locomotion were detected (Fig. 4J), indicating no abnormalities of explorative behavior.

**Figure 4.**
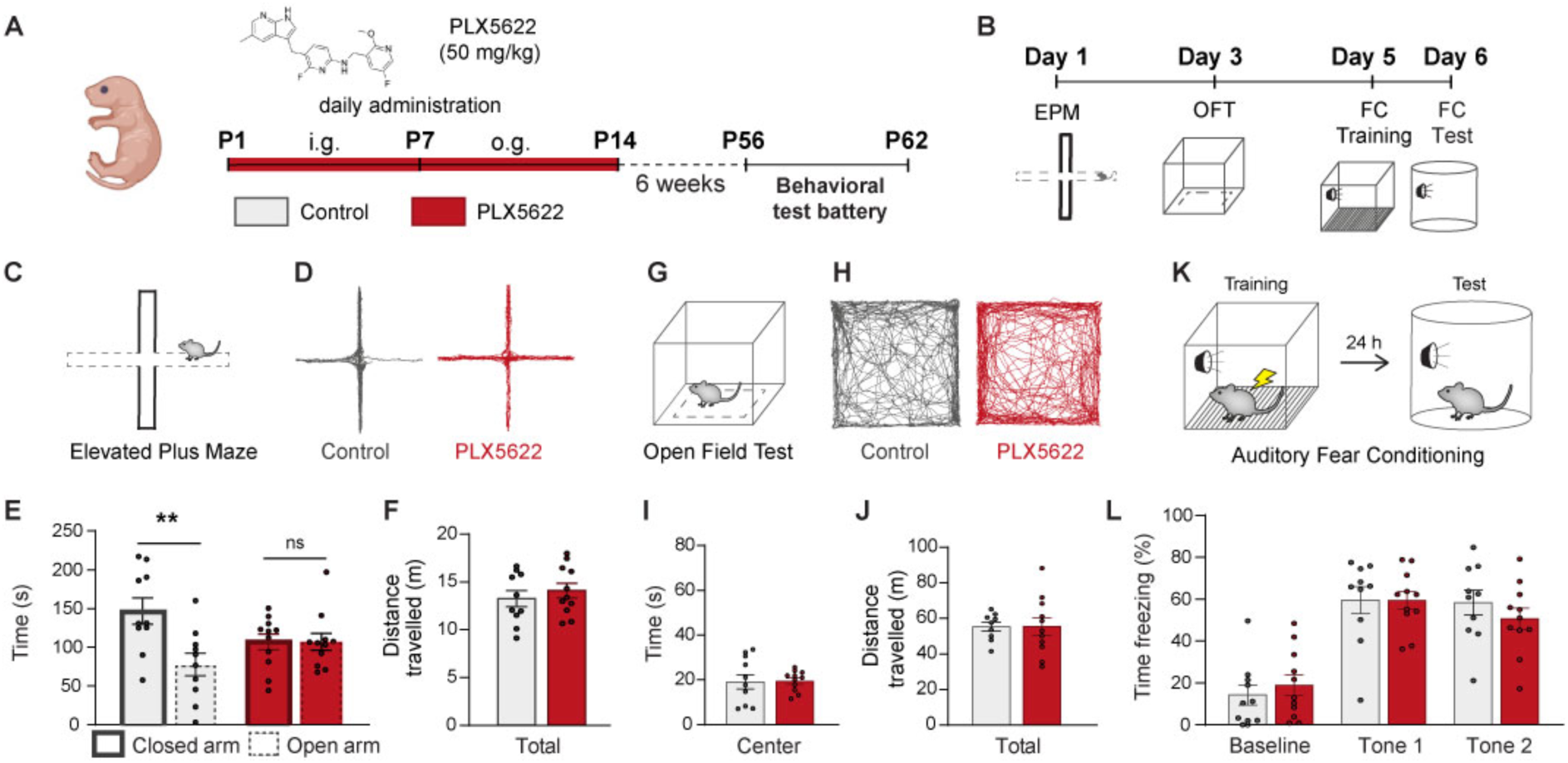
Absence of microglia from P1-P14 impairs anxiety-related behavior on the EPM in adulthood. (**A**) Experimental design for PLX5622-mediated depletion of microglia from P1-P14 and behavioral testing at 8 weeks of age. (**B**) Schematic representation of the order of tests in the behavioral test battery. **(C)** Schematic representation of the EPM. **(D)** Representative EPM traces from control (grey) and PLX5622-treated (red) mice. **(E-F)** Quantification of time spent in the EPM open vs. closed arms (E) and total distance travelled (F). **(G)** Schematic representation of the OFT. (**H)** Representative OFT traces from control (grey) and PLX5622-treated (red) mice. **(I-J)** Quantification of time spent in the OFT center (I) and total distance travelled (J). **(K)** Schematic representation of the auditory fear conditioning test. **(L)** Quantification of percentage of time spent freezing during the test at baseline and during exposure to tones 1 and 2. Control, n = 10; PLX-treated, n = 11. Error bars represent SEM. Statistically significant comparisons are marked with ** p < 0.01.

To complete the behavioral assessment, fear-related learning behavior was tested in the form of an associative fear conditioning test, in which mice learn to associate a previously neutral tone (conditioned stimulus) with an aversive footshock. Memory of this learned fear behavior is assessed 24 h after training by re-exposing mice to the tone and scoring freezing behavior as a measure of fear expression (Fig. 4K). PLX5622-treated mice showed no alteration in freezing compared with control mice during either of two tone presentations on the test day (Fig 4L), indicating that depletion of microglia during synaptogenesis does not impair associative fear conditioning in adulthood.

In conclusion, these findings indicate that depletion of microglia during synaptogenesis (P1-P14) results in disruptions in specific domains of innate anxiety-related behaviors in adulthood, without affecting the acquisition of fear-related memories.

### Absence of microglia during late postnatal development does not impact GABAergic synapses in the corticolimbic regions

In addition to effects on the initial period of synaptogenesis, it has been reported that depletion of microglia may also affect the subsequent period of synaptic refinement and maturation, which includes approximately the third and fourth week of life in mice (*36–38*). To determine whether depletion of microglia during this time period also affects GABAergic synapses in the corticolimbic system, we administered PLX5622 daily to mice from P15-P28 (Fig. 5A, Fig. S1E). We first confirmed that this treatment schedule successfully depleted microglia from the entire mouse brain (Fig. 5B-C, Fig. S1F). Subsequently, we repeated our immunohistochemical characterisation of GABAergic synaptic markers in the ventral hippocampus (SP and SR), amygdala (BLA and CeA) and medial prefrontal cortex (PL II/III and IL II/III) (Fig. 5D). Importantly, no alterations in the growth rate of the pups were noted, in contrast to the effects previously observed during the period of synaptogenesis, hinting at a fundamental developmental role of CSF1R+ cells during the period of synaptogenesis but not during synaptic refinement (Fig. S1G). Moreover, analysis of the number of VIAAT puncta (Fig. 5E-G), gephyrin puncta (Fig. 5H-J), and Nlgn2 puncta (Fig. S3E-H) revealed no difference in any of the brain regions investigated, although a significant increase in the number of GABA_A_Rγ2 puncta was found in the amygdala (Fig. S4A-D). Together, these data demonstrate that, unlike microglial depletion during synaptogenesis, the absence of microglia during synaptic refinement does not lead to major alterations at inhibitory synapses. These findings indicate that microglia play a selective role in shaping the early postnatal development of the corticolimbic GABAergic system during the first two weeks of life, but not during the subsequent refinement of these circuits, and that the time window of synaptogenesis is most vulnerable to microglia-mediated disruption of GABAergic synaptic function.

**Figure 5.**
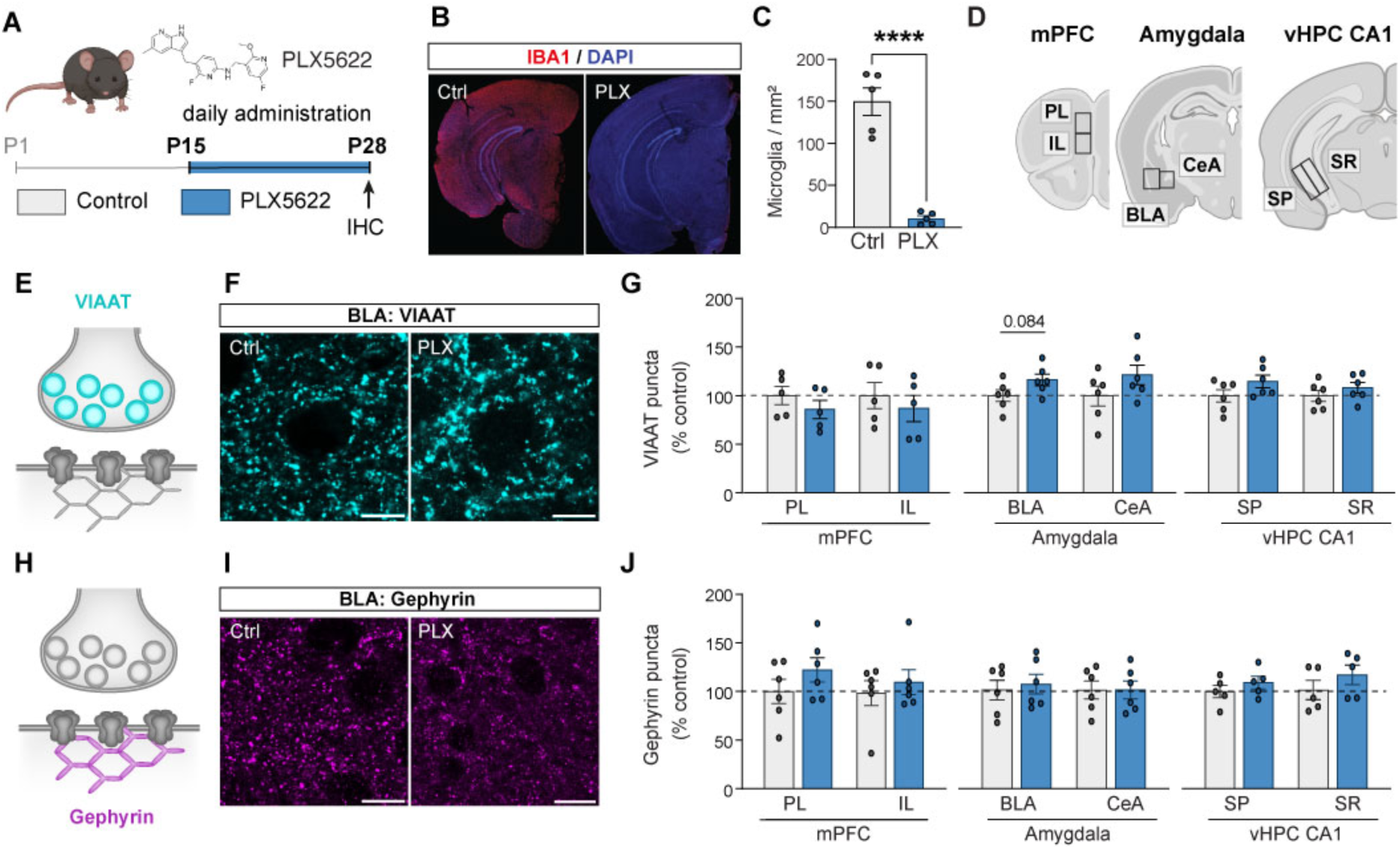
Absence of microglia from P15-P28 does not alter GABAergic synaptic markers in corticolimbic regions. (**A**) Experimental design for PLX5622-mediated depletion of microglia from P15-P28 and immunohistochemical analysis at P28. Image of mouse generated using Biorender. (**B**) Representative photomicrographs of IBA1 immunostaining in control and PLX5622-treated mice. (**C**) Quantification of microglial cell density in control and PLX5622-treated mice (n = 5). (**D**) Anatomical localization of brain regions analysed. **(E)** Synaptic localization of pre-synaptic marker VIAAT. (**F**) Representative photomicrograph of VIAAT immunostaining in control and PLX-treated conditions in the BLA. Scale bar 10 μm. (**G**) Quantification of the number of VIAAT puncta in control (light grey) and PLX5622-treated (blue) mice in each brain region investigated, expressed as percentage of control. (**H**) Synaptic localization of post-synaptic marker gephyrin. (**I**) Representative photomicrograph of gephyrin immunostaining in control and PLX-treated conditions in the BLA. Scale bar 10 μm. (**J**) Quantification of the number of gephyrin puncta in control (light grey) and PLX5622-treated (blue) mice in each brain region investigated, expressed as percentage of control. Error bars represent SEM. Statistically significant comparisons are marked with **** p < 0.0001.

### Absence of microglia during late postnatal development does not lead to altered anxiety-related behaviors in adulthood

To determine whether the lack of alterations in the corticolimbic GABAergic system following microglial depletion during the period of synaptic refinement was accompanied by a similar lack of alterations in anxiety-related behaviors, we next conducted behavioral assessment of these mice in adulthood at 8 weeks of age (Fig. 6A). As described previously, mice were tested on the EPM, OFT and fear conditioning test (Fig. 6B). Consistent with the lack of major synaptic changes, results show no difference between control mice and PLX5622-treated mice in any of the behavioral parameters recorded in the EPM test (Fig. 6C-F), the OFT (Fig. 6G-J) or the fear conditioning test (Fig. 6K-L). These results, together with the data reported in Fig. 5, confirm that microglia do not appear to play a role in shaping GABAergic circuits related to anxiety- and fear-related behaviors during the period of synaptic refinement, supporting the notion of a critical window for microglial function at these synapses in the first two weeks of life in mice.

**Figure 6.**
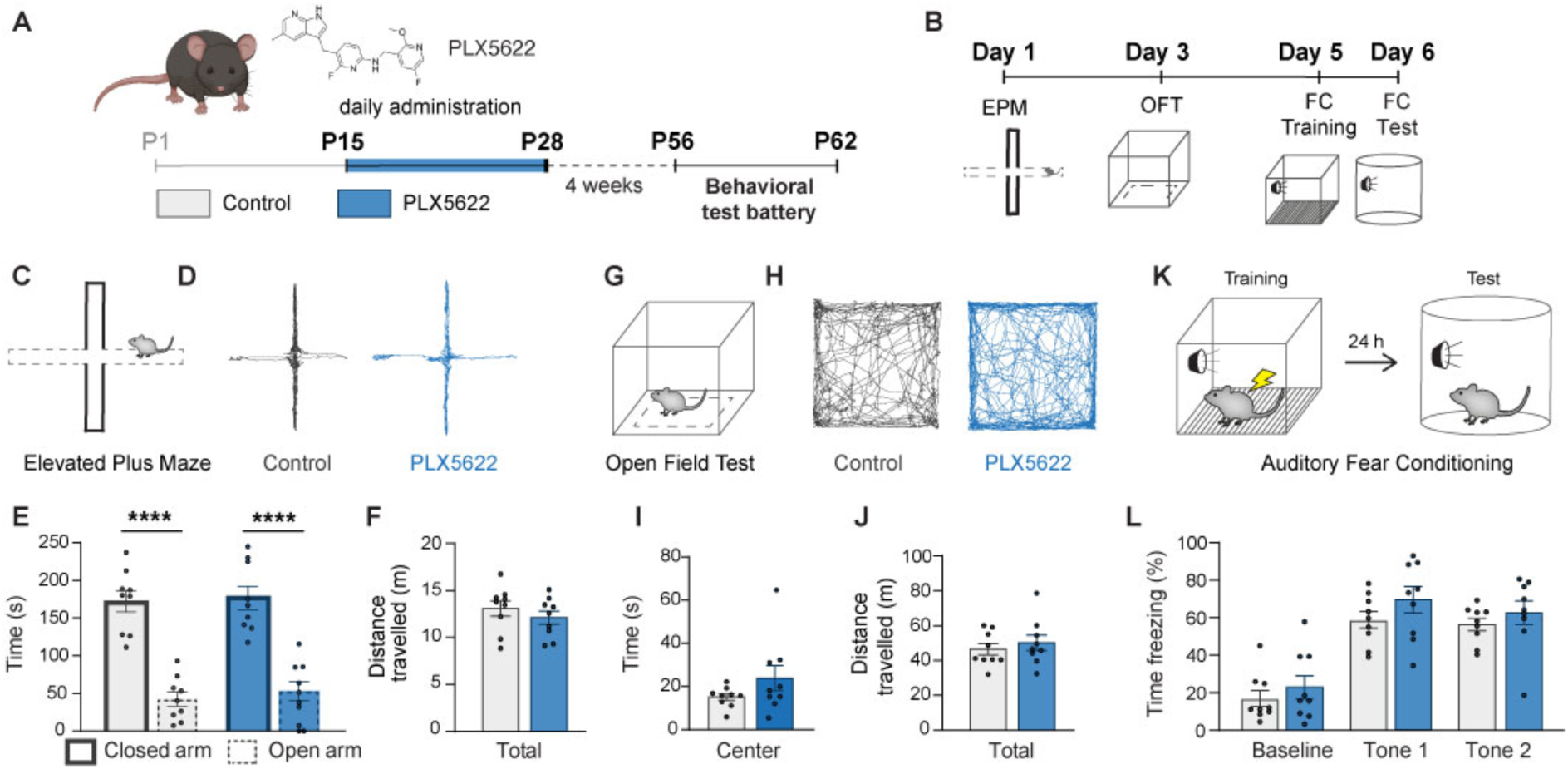
Absence of microglia from P15-P28 does not alter anxiety-related behavior in adulthood. (**A)** Experimental design for PLX5622-mediated depletion of microglia from P15-P28 and behavioral testing at 8 weeks of age. Image of mouse generated using Biorender. **(B)** Schematic representation of the order of tests in the behavioral test battery. **(C)** Schematic representation of the EPM. **(D)** Representative EPM traces from control (grey) and PLX5622-treated (blue) mice. **(E-F)** Quantification of time spent in the EPM open vs. closed arms (E) and total distance travelled (F) (n = 9). **(G)** Schematic representation of the OFT. (**H)** Representative OFT traces from control (grey) and PLX5622-treated (blue) mice. **(I-J)** Quantification of time spent in the OFT center (I) and total distance travelled (J) (n = 9). **(K)** Schematic representation of the auditory fear conditioning test. **(L)** Quantification of percentage of time spent freezing during the test at baseline and during exposure to tones 1 and 2 (n = 9). Error bars represent SEM. Statistically significant comparisons are marked with **** p < 0.0001.

### Loss of GABA_B1_ receptor expression from microglia does not impact corticolimbic GABAergic synapses during early postnatal development

How do microglia detect which GABAergic synapses to remodel? Previous studies in sensory cortical areas have indicated that microglial expression of the metabotropic GABA_B_R may play a key role in detecting GABA release and guiding microglia to the relevant synapses (*25, 26*). To determine whether a similar mechanism may contribute to the observed changes at GABAergic synapses in the BLA and IL at P14, we repeated our immunohistochemical analysis in mice lacking expression of the obligate GABA_B1_R subunit in microglia but not in neurons. To this end, we used mice expressing Cre recombinase under the CX3CR1 promoter (*45*), which selectively labels cells of the macrophage lineage, including microglia, crossed with mice carrying a conditional GABA_B1_R construct (*46*), as described previously (*25, 26*) (Fig. 7A-B, Fig. S4). Analysis of the resulting four genotypes (CX3CR1 wt/wt x GABA_B1_R wt/wt = WT control; CX3CR1 Cre/wt x GABA_B1_R wt/wt = Cre control; CX3CR1 wt/wt x GABA_B1_R fl/fl = loxP-flanked control; CX3CR1 Cre/wt x GABA_B1_R fl/fl = experimental mice) revealed no significant differences between the experimental mice and any of the three controls in the number of VIAAT and gephyrin puncta in the BLA or the IL at P14 (Fig. 7C-J). These data indicate that the previously observed alterations following microglial depletion do not result specifically from the detection of GABA release by microglial GABA_B_ receptors.

**Figure 7.**
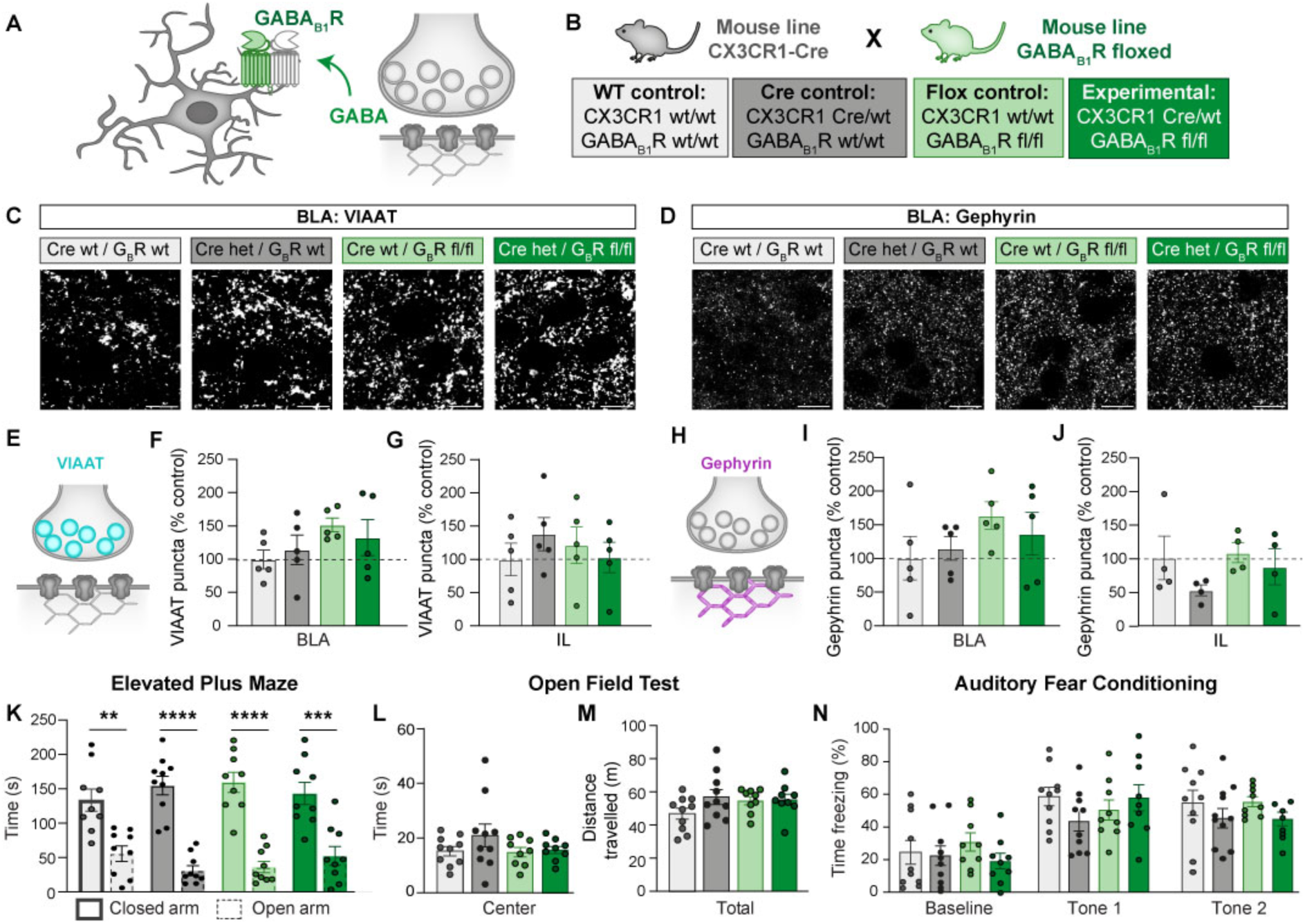
Lack of GABA_B1_ receptor from the microglia does not alter GABAergic synaptic markers at P15 or anxiety-related behavior in adulthood. **(A)** Schematic representation of the previously proposed role for microglial GABA_B1_R in sensing neuronal GABA release (*25*). (**B**) Schematic representation of the genetic strategy used to assess microglial GABA_B1_R function. CX3CR1-Cre and GABA_B1_R fl/fl mouse lines were crossed to generate four genotypes: WT control (light grey), Cre expression control (dark grey), GABAB1R fl/fl control (light green), and experimental mice lacking GABAB1R in CX3CR1-expressing cells (dark green). (**C-D**) Representative photomicrographs of VIAAT (C) and gephyrin (D) immunostaining in the BLA of all four genotypes at P15. (**E**) Synaptic localization of pre-synaptic marker VIAAT. (**F-G**) Quantification of the number of VIAAT puncta in the BLA (F) and IL (G) in all four genotypes at P15, expressed as percentage of WT control. (**H**) Synaptic localization of post-synaptic marker gephyrin. (**I-J**) Quantification of the number of gephyrin puncta in the BLA (I) and IL (J) in all four genotypes at P15, expressed as percentage of WT control. (**K**) Quantification of time spent in the EPM open vs. closed arms by adult mice of all four genotypes (n = 9). (**L-M**) Quantification of time spent in the OFT center (L) and total distance travelled (M) by adult mice of all four genotypes (n = 10,10,9,9). (**N**) Percentage of time spent freezing during the test at baseline and during exposure to tone 1 and tone 2 by adult mice of all four genotypes (n = 10,10,9,9). Error bars represent SEM. Statistically significant comparisons are marked with ** p < 0.01, *** p < 0.001, **** p < 0.0001.

### Loss of GABA_B1_ receptor expression from microglia does not impact anxiety- and fear-related behavior in adult mice

To further determine whether microglial GABA_B_Rs may be required for the disruption in anxiety-related behavior observed in the EPM following microglial depletion in early postnatal development, we repeated the same behavioral test battery (see Fig. 4 and Fig. 6) in CX3CR1-Cre x GABA_B1_R fl/fl mice. No significant differences were observed between the experimental CX3CR1-Cre/wt x GABA_B1_R fl/fl mice compared to any of the three controls in the EPM (Fig. 7K), the OFT (Fig. 7L-M), or in auditory fear conditioning (Fig. 7N). These data demonstrate that the GABA_B1_R expression in microglia is not necessary for adequate behavioral responses in anxiety and fear-related memory tests.

## Discussion

Microglia were recently proposed to play a key role in sculpting GABAergic synapses during physiological brain development, but whether this process contributes to the development of anxiety-related GABAergic circuits in the corticolimbic system remained entirely unknown. In the present study, we report that depletion of microglia from mice during the peak period of synaptogenesis in early postnatal development (P1-P14), but not during the key phases of synaptic refinement in later postnatal development (P15-P28), results in a reduction in the number of perisomatic GABAergic synapses specifically in the amygdala, accompanied by a robust increase in clusters of the GABAergic postsynaptic scaffolding protein gephyrin across several regions of the corticolimbic system. Similarly, early postnatal microglial depletion, but not later postnatal depletion, disrupts the development of normal anxiety-related avoidance behaviors as assessed in the EPM paradigm. Neither of these effects depends on microglial expression of GABA_B1_ receptors, indicating that GABA-receptive microglia do not play a significant role in these processes. Together, our findings lay an important groundwork for understanding the mechanisms by microglia may contribute to the development of anxiety-related GABAergic circuits and their disruption in anxiety disorders.

One of the central objectives of our study was to determine whether microglia play a role in shaping the physiological development of GABAergic synapses in the corticolimbic system. The vast majority of prior studies investigating the interaction between microglia and GABAergic synapses focused on conditions in which the fully developed brain is perturbed, e.g. through lipopolysaccharide administration (*47, 48*), parasite infection (*49*), seizures (*50, 51*), ischemia (*52*), or anesthesia (*53*), or under physiological conditions during learning-related plasticity (*54*). In recent years, a smaller number of investigations have indicated that microglia also contribute to GABAergic synapses in the early development of the cerebellum (*55*) and the somatosensory cortex (*25, 26, 56*). In the present study we extend this analysis to corticolimbic brain regions whose function is particularly vulnerable in psychiatric and neurodevelopmental disorders, including the prefrontal cortex, amygdala, and ventral hippocampus. Our findings indicate that microglia indeed shape GABAergic synapse development in these regions, most prominently in the BLA, providing a basis for future investigations on the mechanisms by which genetically or environmentally induced alterations in this process may contribute to the pathophysiology of psychiatric and neurodevelopmental disorders.

A second key question addressed in our work regards potential time windows of vulnerability for the influence of microglia on synapse development. While the timecourse of synapse formation and refinement is not uniform across brain regions and synapse subtypes, general emerging principles indicate that synapse formation in mice peaks in the first two weeks of life (up to P14-15, corresponding approximately to the first five years of human development), while activity-dependent synaptic refinement and synapse elimination occur predominantly in the second two weeks of life (up to P30, corresponding to later childhood and early adolescence) (*36–38*). Our data indicate that throughout the corticolimbic system, the most pronounced effects of microglial depletion on GABAergic synapse markers are observed during the early postnatal period from P1-P14, with only very modest alterations found in later postnatal development, i.e. from P15-P30. These findings are consistent with previous reports in the somatosensory cortex, where the most vulnerable periods for the remodeling of perisomatic synapses originating from PV-positive basket cells, axo-axonic synapses originating from PV-positive chandelier cells, and dendritic synapses originating from SST-positive cells were identified as P1-P15, P12-P18, and P10-P15, respectively (*25, 26, 56*). Similarly, earlier studies focusing on glutamatergic synapses reported periods of heightened microglial remodeling activity from P8-P14 (*17, 57–59*), although more subtle effects in later developmental time windows were also observed (*60*). These findings indicate that the peak periods of synaptogenesis, rather than later phases of activity-dependent synapse elimination, may be most vulnerable to disruptions in the microglia-dependent development of GABAergic synapses and circuits that contribute to encoding psychiatrically relevant behaviors.

Consistent with this notion, we report that depletion of microglia in early postnatal development (P1-P14), but not in later postnatal development (P15-P28), results in disruptions in anxiety-related avoidance behaviors in adulthood. In the EPM, mice that lacked microglia in this early postnatal period failed to show normal discrimination between the anxiogenic open arms and the ‘safe’ closed arms, instead spending equal amounts of time in both environments. These findings indicate that microglial depletion produces long-lasting disruptions in the normal development of the circuits that encode the innate avoidance of open spaces or heights, which is an adaptive behavior that is essential for the survival of the organism (*8*). It should be noted that at this point, our data do not permit us to causally link the observed alterations at GABAergic synapses in the corticolimbic system with this behavioral impairment. Moreover, anxiety-related avoidance behaviors in the OFT remained unaffected, indicating that not all innate anxiety-related behavioral domains may be disrupted (*61*). Similarly, fear learning in adulthood remained unaffected, likely reflecting the fact that at the time of the induction of learning-related plasticity, microglia had repopulated and were functioning normally. To our knowledge, only one other study that has investigated the consequences of microglial depletion in early postnatal development in mice, reporting that depletion of microglia from P1-P7 using PLX3397 does not alter behavior in the EPM or OFT (*62*). These data would indicate that the disruption in normal anxiety-related avoidance behaviors observed in our study may result specifically from effects in the second postnatal week, broadly consistent with the notion that the most vulnerable period appears to coincide with synapse formation, which peaks around P10-P14 in the brain regions under investigation (*36–38*). In additional support of this notion, numerous studies have observed that microglial depletion in adulthood has no consequences for anxiety-related behaviors under basal conditions, but can influence stress-related behavioral alterations (*63–65*), which may be associated with synaptic rearrangements that re-open a period of heightened vulnerability.

An important area of debate regards the mechanisms by which microglia shape synaptic function. While it was originally proposed that microglia play a key role in pruning excess or inactive synapses via phagocytosis (*17, 40*), more recent studies have indicated that additional mechanisms may contribute to the disruption of synaptic function, include trogocytosis (partial removal of synaptic material), synaptic stripping (physical displacement of synaptic contacts), and destabilization through modulation of the extracellular matrix (*15, 21, 22, 66*). The original concept of pruning has also recently been expanded to distinguish between culling (active removal of synapses from neurons by microglia) and scavenging (uptake of synaptic material previously released by neurons) (*67*). Moreover, microglia have not only been shown to disrupt synaptic function, but conversely also to promote the formation of new synapses or to enhance or protect the function of existing synapses (*20–22, 58, 59*). While the literature specifically relating to GABAergic synapses is substantially more limited, a similar diversity of functions is beginning to emerge (*23, 24*). For example, it has been reported that microglia are required for pruning PV^+^ perisomatic (*25*) and SST^+^ dendritic (*56*) synapses, but promote the development of PV^+^ axo-axonic synapses (*26*), during early postnatal development in the mouse somatosensory cortex. In the present study, we report that microglial depletion in early postnatal development results in region-specific alterations in VIAAT-positive presynaptic terminals, with a slight but significant reduction in VIAAT-positive puncta observed selectively in the BLA. Importantly, the magnitude of this reduction was substantially greater when the analysis was restricted to the perisomatic region, consistent with the notion that it may arise predominantly from an effect on perisomatic PV^+^ basket cell synapses. Moreover, the frequency of mIPSCs was reduced in principal neurons of the BLA. Together, these observations suggest that the number of functional GABAergic synapses is reduced in the BLA following microglial depletion, indicating that microglia likely selectively promote perisomatic synapse formation in this brain region. In contrast, the observed increase in VIAAT puncta in the infralimbic cortex was neither reproduced in the perisomatic-specific analysis nor accompanied by a corresponding change in mIPSCs. It is conceivable that the observed VIAAT increase was driven primarily by effects at distal dendritic synapses, which would likely not be reflected in the mIPSC analysis under our conditions. Nevertheless, the data do not currently warrant the conclusion that GABAergic synapses are upregulated in the mPFC in our study. Intriguingly, however, a robust upregulation was observed throughout the corticolimbic system in the number of clusters of the postsynaptic scaffolding protein gephyrin. This upregulation is likely independent of any changes in synapse number and may result from entirely distinct signaling mechanisms, potentially regulating extrasynaptic accumulations of gephyrin. It has previously been shown that gephyrin distribution can be regulated by the neurotrophic factor BDNF released from microglia (*18, 52*), and that BDNF-mediated regulation of gephyrin in turn contributes to GABAergic signaling in the amygdala (*68*). Whether a similar mechanism is responsible for the gephyrin upregulation observed in our study, and what the functional consequences are for GABAergic synapse function, remains to be determined.

It should be noted that the consequences of microglial function on synapses appear to be influenced on the methodology used to probe microglia. Our investigation, like many other studies focusing specifically on developmental aspects of GABAergic synapses (*25, 26, 56*), employed a pharmacological depletion strategy involving administration of the CSF1R inhibitor PLX5622. Other studies have focused on genetic strategies for microglial depletion or functional modulation (*69, 70*), partially with different results. Most notably, a recent study using the Csf1r^ΔFIRE/ΔFIRE^ model for constitutive microglial depletion reported no significant perturbation of cortical and hippocampal synapses at P14, leading the authors to conclude that microglia are not required for many neurodevelopmental processes (*71*). While this study focused largely on excitatory synapses, spontaneous transmission at GABAergic synapses in the somatosensory cortex was also found to be unaltered. It is conceivable that the discrepancy to prior studies may be due to the fact that the Csf1r^ΔFIRE/ΔFIRE^ study (*71*) did not discriminate between different GABAergic synapse subtypes, which were previously shown to be oppositely regulated (*25, 26, 56*). Indeed, our own research would support this conclusion, since we observed substantially larger effect sizes when selectively analyzing perisomatic GABAergic synapses in the amygdala. It would be interesting to directly address this question in the future by combining synapse subtype-specific analyses with a comparison of pharmacological and genetic manipulation strategies.

How do microglia detect at which GABAergic synapses they should exert their effects? It has been previously shown that certain subtypes of microglia express metabotropic GABA_B_ receptors, and the idea that GABA itself may act as a signaling molecule to attract GABA_B_R-expressing microglia has therefore gained substantial attention (*24, 25, 51, 72, 73*). Consistent with this notion, it was reported that genetic deletion of GABA_B_R1 from CX3CR1-positive cells, including microglia, abolishes the function of these microglia at GABAergic synapses in the somatosensory cortex (*25, 26*). In contrast, our findings demonstrating that microglial effects on GABAergic synaptic puncta in the BLA and mPFC and on anxiety-related avoidance behavior in the EPM are independent of microglial GABA_B_R1 expression (Fig. 7), indicating that other signaling pathways must recruit microglia to the relevant GABAergic synapses. While a substantial number of potential signaling pathways have been identified that link microglia to glutamatergic synapses (*18, 20, 74, 75*), less is known about their specific role in the recruitment to GABAergic synapses (*24*). The display of the “eat me” signal phosphatidylserine on GABAergic synapses was reported to lead to MERTK-dependent microglial recruitment and pruning of the respective GABAergic synapses (*76*). Signaling through ATP binding to purinergic receptors on microglia (particularly P2Y_12_ or P2RX7 receptors) has also been implicated (*50, 77*), and a recent study indicates that this may involve astrocyte-to-microglia purinergic signaling (*78*). Little evidence is currently available for the involvement of other classical microglia-synapse interaction pathways at developing GABAergic synapses, such as the complement or fractalkine pathways. Our data indicate that a broader analysis beyond the current focus on GABA-receptive microglia will be essential for fully understanding the role of microglial remodeling of GABAergic synapses in psychiatrically relevant circuits.

## Materials and methods

### Experimental subjects

Mice were maintained at the Translational Animal Research Center of the Johannes Gutenberg University Mainz on a 12h light/dark cycle with water and food ad libitum. Only male mice were used for experiments. For experiments involving pharmacological depletion of microglia via PLX5622 administration (Fig. 1–6), pregnant C57BL/6JRj females were obtained from Janvier at E13 and housed until birth of the pups. Male experimental pups were maintained with their mothers until weaning at P21, and PLX5622 administration occurred at P1-14 or P15-28 as described below. For experiments involving genetic deletion of GABABRs from microglia (Fig. 7), loxP-flanked GABA_B1_R mice ((*46*), generously provided by Dr. Bernhard Bettler, University of Basel) and CX3CR1-Cre mice ((*45*), generously provided by Dr. Steffen Jung, Weizmann Institute) were backcrossed for 5-6 generations onto a C57BL/6JRj background. The two mouse lines were subsequently crossed to generate GABAB1R fl/fl x CX3CR1-Cre heterozygous breeders. To generate experimental mice, GABAB1R fl/fl x CX3CR1-Cre heterozygous male breeders were crossed with GABAB1R fl/fl female breeders. This breeding strategy generated experimental male littermates with the following genotypes: (1) GABAB1R wt/wt x CX3CR1-Cre wt (= WT control), (2) GABAB1R wt/wt x CX3CR1-Cre het (= Cre control), (3) GABAB1R fl/fl x CX3CR1-Cre wt (= loxP-flanked control), and (4) GABAB1R fl/fl x CX3CR1-Cre het (experimental group).

For both pharmacological and genetic manipulation experiments, immunohistochemical analysis was conducted at P14-P16, P28-P30, or 8-10 weeks, as described below. Electrophysiological analysis was conducted at P14-P16. Behavioral experiments were conducted at 8-10 weeks. Experimenters were blind to the experimental condition and genotype throughout data acquisition and analysis; unblinding occurred once all data had been acquired and analysed. All procedures were conducted in accordance with the guidelines of the welfare of experimental animal use issued by the federal government of Germany and the Johannes Gutenberg University Mainz and were approved by the Landesuntersuchungsamt Rheinland-Pfalz under the license G 22-1-034.

### PLX5622 preparation and administration

Microglial depletion from P1-P14 was achieved through administration of PLX5622 at a final concentration of 50 mg/kg in diluent solution, which consisted of hydroxypropyl methylcellulose (HPMC, Sigma Aldrich, #H7509-25G) 0.5% and polysorbate 80 (PS80, Sigma Aldrich, #W291706-1K-K) 1% in Type I ultrapure water (Sartorius Arium Mini System). To prepare the final working solution, PLX5622 powder (Med Chem Express, #HY-114153) was initially diluted in dimethyl sulfoxide (DMSO, Carl Roth, #4720.1) at a concentration of 0.25 mg/μl and subsequently aliquoted with further dilution steps with DMSO to obtain the correct working amount of PLX5622 per injection, adjusted to the estimated body weight for each injection day. Aliquots of PLX5622 stock solution were stored at -80°C for a maximum of 6 months and thawed immediately prior to initiation of experimental procedures. Diluent solution was freshly prepared every experimental week from stock solutions (stock of HPMC, 2% in ultrapure water; stock of PS80, 25% in ultrapure water) and was stored at room temperature (RT) in the dark. PLX5622 working solution (consisting of 95% diluent solution and 5% PLX5622 stock solution) was prepared freshly for each administration day, using bodyweight-adjusted aliquots as described above. For control animals, control solution (consisting of 95% diluent solution and 5% DMSO) was prepared. Before each injection, the working solution was treated in a warm bath for 1h followed by homogenization with the Bandelin SONOPLUS mini20 for 90 sec with an amplitude of 25%. Working solution was administered to the mice within 10 min after homogenization to prevent reprecipitation of PLX5622. For mice at P1-P7, intragastric injections were performed, while for mice at P8-P20 the working solutions were administered with oral gavage. Drug administration was conducted daily between 12 and 2 pm. PLX5622 administration at P21-P28 was provided through PLX5622-containing chow (ssniff Spezialdiäten GmbH, 1200 ppm PLX5622, available ad libitum). Control animals received the same chow without PLX5622.

### Immunohistochemistry procedures

For all immunohistochemistry procedures, mice were anesthetized with isoflurane and subsequently sacrificed by decapitation. The brains were removed and split into two hemispheres. One hemisphere was fixed in 4% paraformaldehyde (PFA) (Carl Roth, P087.1) overnight (O/N) at 4°C for analysis of microglia depletion, then transferred to 30% sucrose (Sigma Aldrich, #S0389-5KG) in ultrapure water at 4°C for cryoprotection. The other hemisphere was fresh-frozen in 2-methylbutane at -35°C for approximately 30 seconds for synaptic analysis and then stored at -80°C.

To confirm effectiveness of microglial depletion via IBA1 immunohistochemistry, 40 μm thick brain sections were prepared from the PFA-fixed hemispheres using a cryostat (Leica, #CM1950, specimen temperature -20°C, chamber temperature -17/18°C) and stored in phosphate buffered saline (PBS) containing 0.1% sodium azide. Sections were washed 3x 10 min in PBS, followed by incubation in blocking solution (10% NGS, Merck, #S26-100ML; 0.3% Triton X-100, Sigma Aldrich, #X100-1L; in PBS) for 1h at RT. Subsequently, sections were incubated O/N at 4°C with primary antibody (anti-IBA1 antibody, RRID: AB_839504, Wako, 019-19741, 1:1000) diluted in 8% NGS and 0.3% Triton-X-100 in PBS. Sections were then washed 3x 10 min with PBS, then incubated with secondary antibody (goat anti-rabbit Alexa Fluor (AF) 647, AB_2535813, Thermo Fisher, #A-21245, 1:600) and 4′,6-diamidino-2-phenylindole (DAPI, ThermoFisher Scientific, #D1306, 1:5000) for 2h at RT. Sections were washed 3x 10 min in PBS at RT, then mounted onto microscopy slides and coverslipped with mounting medium (ROTI Mount FluorCare, Carl Roth, No. HP19.1).

To quantify markers of GABAergic synapses, 18 μm thick brain sections were prepared from the fresh-frozen hemispheres using a cryostat (Leica, #CM1950, specimen temperature - 20°C, chamber temperature -17/18°C) and mounted directly onto microscopy slides. Sections for matched sets of brains were mounted on the same slide to ensure identical staining conditions between genotypes and experimental conditions (i.e. sections from control and PLX5622-treated brains for pharmacological microglia depletion experiment; sections from WT control, CX3CX1-Cre control, and loxP-flanked control brains as well as brains of experimental mice for genetic microglial manipulation experiment). After mounting, brain sections were dried at RT for 30 minutes. They were then fixed using one of two fixation methods, i.e. methanol fixation or PFA post-fixation, depending on the primary antibodies used as specified below. 1) For methanol fixation, brain sections were fixed in methanol at -18°C for 5 min. 2) For PFA post-fixation, sections were fixed in 4% PFA at RT for 5 min, followed by epitope retrieval in pre-heated citric buffer (0.01M in dH2O, pH 6) at 100°C for 30 min and subsequent cooling in the same solution at RT for 15 min. After fixation, sections were washed 3x 10 min in PBS and incubated for 1h at RT with blocking solution (10% NGS and 0.3% Triton X-100 in PBS) for 1h at RT in a humid chamber. After blocking, sections were incubated with primary antibodies diluted in 8% NGS, 0.3% Triton X-100 in PBS O/N at 4°C. The following primary antibodies were used: 1) Methanol fixation: guinea pig anti-Nlgn2 (RRID:AB_2619815, SySy #129205, 1:500), mouse anti-Gephyrin (RRID:AB_2619837, SySy #147111, 1:500). 2) PFA post-fixation: mouse anti-VIAAT (RRID:AB_2619818, SySy #131011, 1:1000), rabbit anti-GABA_A_Rγ2 (RRID:AB_2263066, SySy #224003, 1:1000), rat anti CD68 (RRID:AB_2044003, BioLegend #137001, 1:1000). On the next day, sections were washed 3x 10 min in PBS, then incubated for 2h at RT with secondary antibodies and DAPI (1:5000) in 8% NGS, 0.3% Triton X-100 in PBS, and again washed 3x 10 min in PBS. The following secondary antibodies were used (all at 1:600, all from Thermo Fisher): Goat anti-rabbit AF488 (RRID:AB_2866491, #A32731TR), goat anti-mouse AF555 (RRID:AB_2535844, #A-21422), goat anti-guinea pig AF488 (RRID:AB_2534117, #A-11073). Once the washes were completed, sections were coverslipped with mounting medium ROTI Mount FluorCare (Carl Roth, No. HP19.1).

### Image acquisition and processing for analysis of microglial depletion

Images were acquired using a Leica TCS-SP8 laser scanning confocal microscope (Leica microsystems, Germany) equipped with a white light laser as well as Hybrid Detector (HyD) and photomultiplier tube (PMT) detectors for signal processing. A 20x objective with a 0.75x digital zoom factor was used to acquire tiled single plane images of coronal brain sections at 512 x 512 spatial resolution for both DAPI and AF647. and images were stitched for smoothening of the borders of single tiles. Laser power was optimized for a dynamic range of detection that excluded oversaturation of both DAPI and AF647 signals. All imaging parameters were kept constant for images acquired from matched pairs of control and PLX5622-treated brains.

Image analysis was conducted using the Fiji ImageJ software (*79*). Images were first binarized using a constant threshold for all sections. Microglial cell bodies were then counted using the function “Analyze Particles” with the size range of 50 μm^2^ -infinity, with no circularity filters, in order to include microglial cells of any shape. The number of cells was divided by the area of the coronal section, in order to determine the number of microglial cells per mm^2^.

### Image acquisition and processing for synaptic analysis

Images were acquired using a Leica TCS-SP8 laser scanning confocal microscope equipped as described above. A 63x objective with a 4x digital zoom factor was used to acquire single plane images with a 1024 x 1024 spatial resolution. Ten images were acquired within the same imaging session from each experimental condition (control vs PLX5622-treated for pharmacological experiment; WT control, Cre control, loxP-flanked control and experimental sample for the genetic experiment) for each of the six brain regions analysed: Stratum pyramidale (SP), Stratum radiatum (SR), basolateral amygdala (BLA), central amygdala (CeA), prelimbic cortex layer II/III (PL) and infralimbic cortex layer II/III (IL). Laser power was optimized for a dynamic range of detection that excluded oversaturation of the wavelength used to detect the synaptic signal. All imaging parameters were kept constant for images acquired from matched sets of experimental conditions within each imaging session.

Image analysis was conducted using the Fiji ImageJ software (*79*). Images were first binarized using an identical threshold for each experimental condition to be compared (see above). To determine the threshold value, the intersynaptic background intensity was measured from all images acquired from one brain region within one session, and the mean background intensity was calculated across all images and multiplied by a factor of 2.5 (i.e. threshold = 2.5 x (sum of intersynaptic background from 10 images from each experimental condition) / (10 images x number of conditions)). After binarization, images were subjected to ‘noise despeckle’ and ‘watershed segmentation’ function. The resulting images were then subjected to the “analyze particles” function using a filter of 0.03-1 μm^2^ for gephyrin, Nlgn2 and GABA_A_Rγ2 puncta and 0.03-3 μm^2^ for VIAAT puncta. Number and size of puncta were quantified for each image, normalized to the average value from all images within the experimental condition and expressed as % of control condition for each experiment.

### RNA in situ hybridization analysis of GABA_B1_R expression in microglia

To confirm cell type-specific deletion of GABA_B1_R expression from CX3CR1^+^ cells, RNA in situ hybridization was performed using the RNAscope Multiplex Fluorescent V2 Assay kit (Advanced Cell Diagnostics, Newark, CA, USA) according to the manufacturer’s instructions. Probe detection was conducted using Opal520 (Akoya Biosciences #SKU FP1487001KT). Co-localization with microglia was assessed using immunohistochemical detection of IBA1 (anti-IBA1 antibody, RRID: AB_839504, Wako, 019-19741, 1:1000) as described above.

### Image acquisition for analysis of GABA_B1_R expression in microglia

Images were taken using a confocal microscope Leica SP8 (Leica microsystems, Germany) equipped with HyD and PMT detector for signal processing. Using a 40x objective with a 2x zoom factor, with a 1024×1024 spatial resolution, single microglial cells where individuated. GABBR1 mRNAs were detected with Opal 520 and IBA1, microglial marker, with AF647. Each cell was captured with Z-stack imaging with each Z-step size equal to 0.3 μm. Lasers were optimized in power for a dynamic range of detection that excluded oversaturation of both Opal 520 and AF647 signals; imaging settings were kept constant for each genetic condition. Image analysis was conducted using the IMARIS program. Colocalization of GABBR1 mRNAs expression with microglia was analysed using 3D reconstruction of Z-stack microglial images.

### Electrophysiological recordings

P15 (+/- 1 day) male mice were anesthetized using isoflurane and subsequently decapitated for brain dissection in ice-cold NMDG solution (in mM: 93 NMDG, 2.5 KCl, 1.2 NaH2PO4, 30 NaHCO3, 20 HEPES, 25 Glucose, 5 L-Ascorbic Acid, 3 Myo-Inositol, 3 Na-Pyruvate, 93 HCl, 10 MgCl2, 0.5 CaCl2; pH adjusted to ∼7.4 with HCl, 300 ± 10 mOsm). Coronal sections containing the mPFC or the amygdala were prepared at 270 μm thickness using a vibratome (Leica, #VT1200). During sectioning, the vibratome chamber was filled with ice-cold NMDG solution, constantly carbonated with 95% O_2_ and 5% CO_2_. After slicing, brain sections were placed for recovery in aCSF solution (in mM: 125 NaCl, 2.5 KCl, 25 NaHCO_3_, 1.25 NaH_2_PO_4_, 10 Glucose, 2 sodium-pyruvate, 3 myo–inositol, 0.44 ascorbic acid, 2 CaCl_2_ and 1 MgCl_2_; pH = 7.3 adjusted by bubbling with carbogen, ∼300 mOsm) at 35°C for 20 minutes and subsequently maintained in aCSF at RT for up to 8h. Patch pipettes (3–5 MΩ open tip resistance) were pulled from borosilicate glass (Science Products, #GB150(F)-8P; Puller: Sutter Instruments, #P-1000). Neurons in the regions of interest were visually identified using an upright microscope (Scientifica, SciCam+ Camera) equipped with infrared detection and 4x and 60× objectives (Olympus). Whole-cell patch-clamp recordings were conducted at RT. During recordings, slices were continuously perfused with carbonated aCSF at a rate of 1.5–2ml/min. The holding potential was set to −70 mV and miniature inhibitory postsynaptic currents (mIPSCs) were recorded under voltage clamp with patch pipettes filled with internal solution (in mM: 54mM KCl, 4 NaCl, 56mM K-gluconate, 20mM HEPES, 0.1mM EGTA, 10 mM Phosphocreatine, 2mM MgATP, 0.3mM Na-GTP; pH adjusted to ∼7.3 with KOH and filtered twice before usage, ∼290 mOsm). To pharmacologically isolate GABAergic postsynaptic currents, 10 μM 6-cyano-7-nitroquinoxaline-2,3-dione (NBQX) (HelloBio, HB0443), 10 μM (R)-3-(2-Carboxypiperazin-4-yl)-propyl-1-phosphonic acid ((R,S)-CPP) (HelloBio, HB0036) and 1 μM tetrodotoxin citrate (TTX) (HelloBio, HB1035) were added to the aCSF bath to block AMPARs, NMDARs, and to block voltage-activated Na^+^ currents and suppress action potential firing respectively. Recordings with an initial series resistance of >20 MΩ or a leak current >300 pA were discarded. Patch-clamp data were acquired with an Axon Digidata 1550 B Data acquisition system (Molecular Devices) with a MultiClamp 700B amplifier (Molecular Devices) and the Clampex Software (PClamp 11) using a low-pass Bessel filter with a cut-off frequency of 10 kHz and digitized at 20 kHz. All offline analyses were performed with IgorPro (Wavemetric, USA). Membrane resistance and cell capacitances were estimated from current transients recorded under voltage clamp in response to 10 mV depolarizing voltage steps from a holding potential of −70 mV and calculated by assuming an interpreted two parts compartment equivalent circuit model. mIPSCs were detected using a sliding template-matching algorithm implemented in IgorPro (*80*) after additional offline filtering using a Gaussian low-pass filter with a cut-off frequency of 2 kHz. Amplitudes as well as rise and decay time of individual mIPSCs were analysed as previously described (*81*).

### Elevated Plus Maze test (EPM)

EPM testing was conducted in a plus-shaped metal apparatus with two open arms and two enclosed arms (arm length 27 cm, arm width 7 cm, wall height 40 cm, height from ground 42 cm, light intensity 60 lux). Each mouse was placed into the center of the apparatus at the intersection of the two arms, and explorative behavior was recorded with an overhead camera for 5 min. Performance was scored using the EthoVision XT software (Noldus). Between each mouse, the apparatus was cleaned with ultrapure water and 70% EtOH to avoid interference of odours between tests. Parameters analysed included total distance travelled and time spent in each arm.

### Open Field test (OFT)

OFT testing was conducted in a square metal apparatus (50 x 50 x 50 cm, light intensity 60 lux. Each mouse was placed into the corner of the apparatus to start the experiment, and explorative behavior was recorded with an overhead camera for 10 min. Performance was scored using the EthoVision XT software (Noldus). Between each mouse, the apparatus was cleaned with ultrapure water and 70% EtOH. Parameters analysed included total distance travelled, average speed, number of entries into the center and time spent in the inner center (15 x 15 cm) and periphery (outer 10 cm along each side of the apparatus).

### Auditory Fear Conditioning Test

Auditory fear conditioning was conducted using a near infrared (NIR) video fear conditioning system (MedAssociates. Fairfax, VT, USA), consisting of two fear conditioning chambers (30 x 24 x 21 cm) with distinct internal design to provide two distinguishable contexts, context A and context B. In context A, the floor consisted of a metal grid capable of delivering an aversive foot shock, and the walls consisted of metal plates. In context B, both the walls and the floor consisted of white plastic plates. Fear conditioning was conducted across two days, including a training day (pairing of a neutral auditory stimulus with an aversive foot shock in context A) and a test day (retrieval of the memory of this association in context B). Freezing was scored throughout both days as a measure of fear-related defensive behavior using the VideoFreeze software.

On the training day (day 1), mice were placed into the first fear conditioning chamber (context A) and allowed to explore the chamber for 2 min. During this time, baseline freezing levels were recorded. Subsequently, a neutral tone (30 s, 80 dB) was presented, co-terminating with a foot shock (2 s, 0.5 mA) delivered through the metal grid floor. After an inter-trial interval (ITI) of 60 s, the sound exposure and co-terminating footshock were repeated. 60 seconds after the second shock, the experimental mouse was placed in a separate resting cage until all the cagemates were tested, before being returned to the homecage, in order to avoid post-test influences on the cagemates. Between each test, the chamber was cleaned first with water and then with 1% acetate.

On the test day (day 2), mice were placed into the second chamber (context B) and allowed to explore the chamber for 2 min, during which baseline freezing levels were recorded. Subsequently, the same tone (30 s, 80 dB) as on day 1 was presented, but without the footshock. After an ITI of 60 s, the sound exposure was repeated. Subsequently, the experimental mouse was placed in a separate resting cage before being returned to the homecage, as described above. Between each test, the chamber was cleaned first with water and then with 70% EtOH.

### Statistical analysis

Statistical analysis was performed using Prism (GraphPad Software, La Jolla, CA, USA). Each data set was initially tested for normality using the Shapiro-Wilk test. In addition, outliers were detected using the ROUT test (1%) and removed from the data sets. For the timecourse of microglial interaction with GABAergic presynapses, a one-way ANOVA with Tukey’s multiple comparison test for post-hoc analysis was conducted. For experiments involving the pharmacological depletion of microglia, PLX5622-treated vs control groups were analysed using an unpaired, two-tailed Student’s t-test (when normally distributed) or a Mann-Whitney test (when not normally distributed). Exception was made for the behavioral experiment EPM and FC where a two-way ANOVA with Tukey’s multiple comparison test for post-hoc analysis was conducted, with the “PLX vs Control” factor and the “Open Arms vs Closed Arms” factor for the EPM, and “PLX vs Control” factor and the “FC Phase” for the fear conditioniong test. To assess the effect of PLX5622 on the body weight of the mice during P1-P15 and P15-P28, a two-way ANOVA Tukey’s multiple comparison test was conducted. Data sets from behavioral analysis of CX3CR1-Cre x GABA_B1_R fl/fl mice and littermate controls were subjected to two-way ANOVA Tukey’s multiple comparisons test for post-hoc analysis (EPM), one-way ANOVA Tukey’s multiple comparisons test (OFT) and to two-way ANOVA Sidack’s multiple comparison test (fear conditioning).

## Acknowledgements

This work was supported by the Deutsche Forschungsgemeinschaft (DFG), Heisenberg grant KR5239/1 (to D.K.-B.) and SPP 2395 (project number 500247271, grant KR5239/3 to D.K.-B. and WA 1600/17-1&2 to A.W.). It was further supported by the Forschungsinitiative Rheinland-Pfalz through the ReALity Excellence Program (D.K.-B. and A.W.) and by the Werner Reichenberger Foundation (M.J.S.). The authors thank Dr. Bernhard Bettler, University of Basel, for providing the GABA_B1_R loxP-flanked mouse line, and Dr. Steffen Jung, Weizmann Institute of Science for providing the CX3CR1-Cre mouse line. The authors are grateful to Marlena Blöchle, Mikkel Blake, Alexander Wenzel, Christine Oswald and the staff at the Translational Animal Research Center for excellent technical support and animal care.

## Author contributions

D.K.-B. conceived and supervised the study, and D.K.-B. and F.R. designed experiments. F.R. performed PLX5622 administration for all experiments, as well as immunohistochemical and behavioral analysis of consequences of PLX5622 administration and microglial GABA_B1_R deletion. K.S. and T.R. performed and analysed electrophysiological recordings. J.S. performed immunohistochemical analysis of microglial engulfment of GABAergic synapses. M.J.S. and A.W. provided conceptual advice, supervision, and experimental materials. D.K.-B. and F.R. wrote the manuscript, F.R., K.S., T.R. and J.S. prepared the figures, and all authors edited and approved the final manuscript.

## Data and materials availability

All data needed to evaluate the conclusions of the paper are present in the paper and/or the Supplementary Materials or are available from the corresponding author upon reasonable request.

