## Supplementary Material for "Microglia regulate early postnatal development of corticolimbic GABAergic synapses and anxiety-related behavior"

**Supplementary Figure 1**

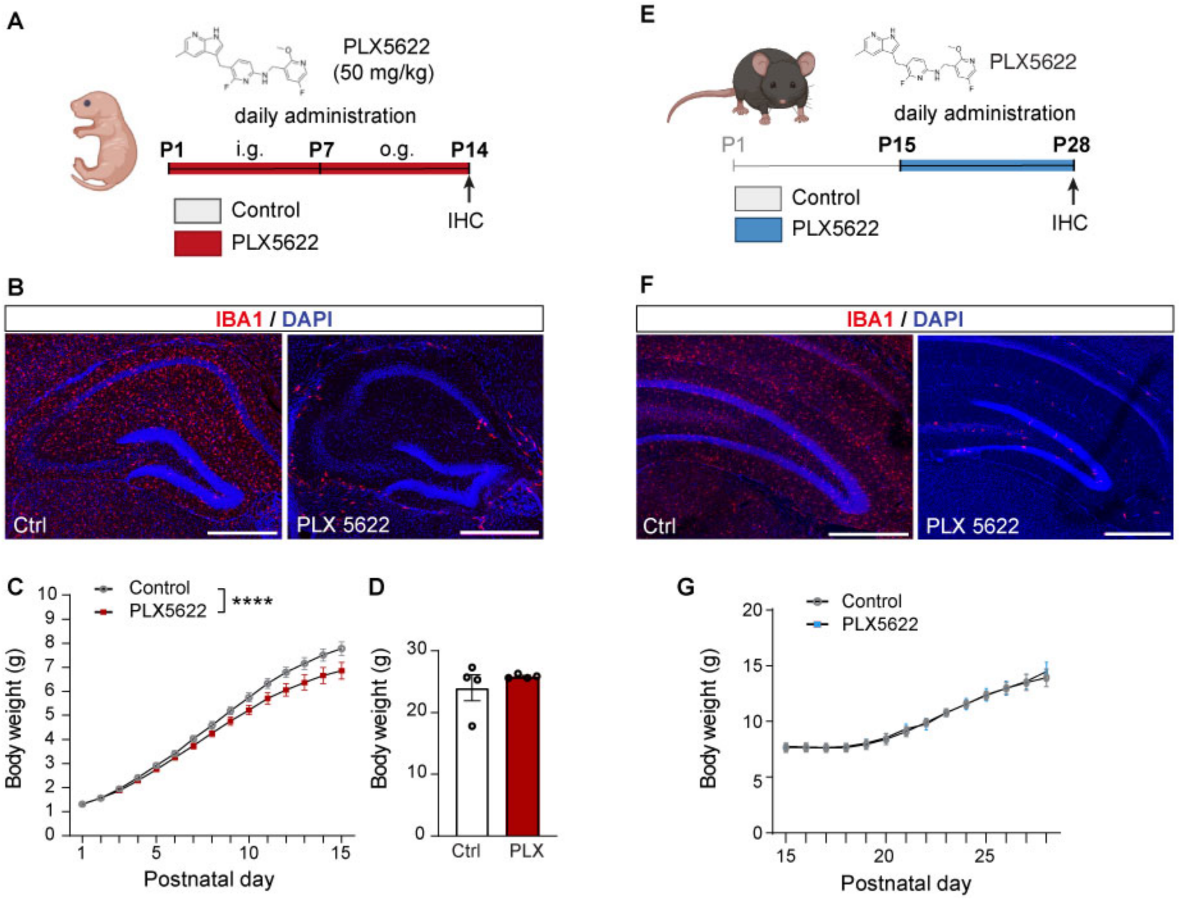


**Fig. S1. Effects of PLX5622 administration on microglial depletion in the hippocampus and on body weight. (A)** Experimental design for PLX5622-mediated depletion of microglia from P1-P14 and immunohistochemical analysis at P14. Image of mouse generated using Biorender. (**B**) Representative photomicrographs of Iba1 immunostaining in control and PLX5622-treated mice at P14. (**C**) Quantification of body weight of control and PLX5622-treated pups from P1-P14 (n = 12). **(D)** Quantification of adult body weight (at 8 weeks of age) of mice that had received control or PLX5622 treatment from P1-P14. **(E)** Experimental design for PLX5622-mediated depletion of microglia from P15-P28 and immunohistochemical analysis at P28. Image of mouse generated using Biorender. (**F**) Representative photomicrographs of Iba1 immunostaining in control and PLX5622-treated mice at P28. **(G)** Quantification of body weight of control and PLX5622-treated pups from P1-P14 ( (n = 6). Error bars represent SEM. Statistically significant comparisons are marked with **** $p < 0.0001$.

### Supplementary Figure 2

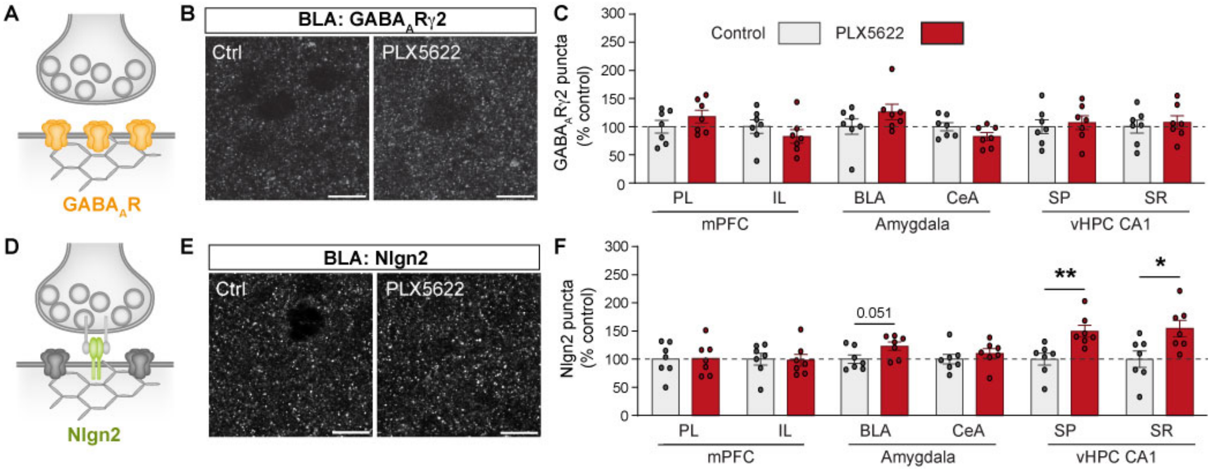


**Fig. S2. Effect of absence of microglia from P1-P14 on GABA$_A$Rγ2 and Nlgn2 at GABAergic synapses in the corticolimbic system. (A)** Synaptic localization of post-synaptic GABA$_A$Rs. (**B**) Representative photomicrograph of GABA$_A$Rγ2 immunostaining in control and PLX5622-treated conditions in the BLA. Scale bar 10 µm. (**C**) Quantification of the number of GABA$_A$Rγ2 puncta in control (light grey) and PLX5622-treated (red) mice in each brain region investigated, expressed as percentage of control (n = 7). (**D**) Synaptic localization of post-synaptic Nlgn2. (**E**) Representative photomicrograph of Nlgn2 immunostaining in control and PLX-treated conditions in the BLA (n = 7). Scale bar 10 µm. (**F**) Quantification of the number of Nlgn2 puncta in control (light grey) and PLX5622-treated (red) mice in each brain region investigated, expressed as percentage of control. Error bars represent SEM. Statistically significant comparisons are marked with * $p < 0.05$, ** $p < 0.01$.

### Supplementary Figure 3

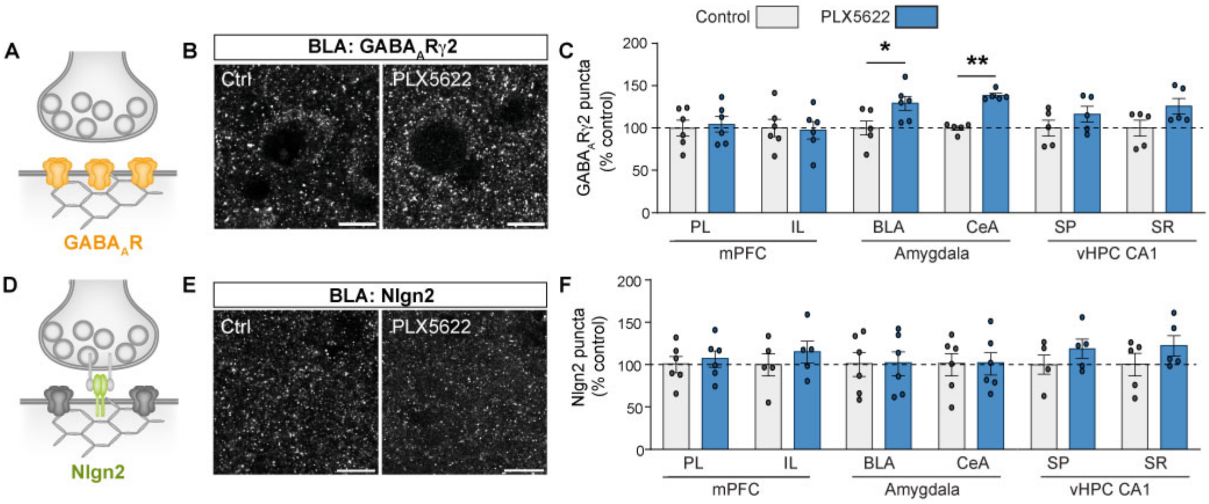


**Fig. S3. Effect of absence of microglia from P15-P28 on GABA$_A$Rγ2 and Nlgn2 at GABAergic synapses in the corticolimbic system. (A)** Synaptic localization of post-synaptic GABA$_A$Rs. (**B**) Representative photomicrograph of GABA$_A$Rγ2 immunostaining in control and PLX5622-treated conditions in the BLA. Scale bar 10 µm. (**C**) Quantification of the number of GABA$_A$Rγ2 puncta in control (light grey) and PLX5622-treated (blue) mice in each brain region investigated, expressed as percentage of control (n = 5-6). (**D**) Synaptic localization of post-synaptic Nlgn2. (**E**) Representative photomicrograph of Nlgn2 immunostaining in control and PLX-treated conditions in the BLA (n = 5-6). Scale bar 10 µm. (**F**) Quantification of the number of Nlgn2 puncta in control (light grey) and PLX5622-treated (blue) mice in each brain region investigated, expressed as percentage of control. Error bars represent SEM. Statistically significant comparisons are marked with * $p < 0.05$, ** $p < 0.01$.

### Supplementary Figure 4

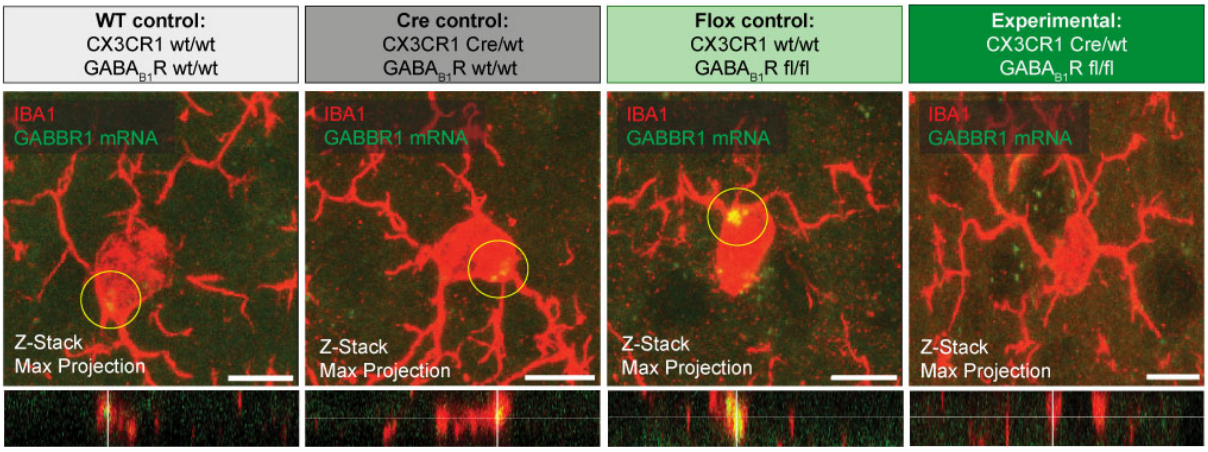


**Fig. S4. Validation of deletion of GABAB1R expression in microglia in CX3CR1 Cre x GABA$_{B1}$R fl/fl mice.** RNAscope detection of GABBR1 mRNA (green) in IBA1-positive cells (red) in all four experimental genotypes as defined in Figure 5. Bottom row: XZ orthogonal view of GABBR1 mRNA in microglia.
